# The fusion peptide proximal region of the HIV-1 envelope glycoproteins regulates exposure of immunogenic epitopes at the trimer base

**DOI:** 10.1101/2020.10.28.360131

**Authors:** Roberth Anthony Rojas Chávez, Devlin Boyt, Changze Han, Li Wu, Hillel Haim

**Affiliations:** Department of Microbiology and Immunology, Carver College of Medicine, The University of Iowa, Iowa City, Iowa, USA

**Keywords:** HIV-1, envelope glycoproteins, fusion peptide proximal region, membrane proximal external region, population-level evolution, antibody neutralization, immune selection

## Abstract

The error-prone replication machinery of HIV-1 continuously generates new variants of the envelope glycoproteins (Envs). Antibody selection pressures applied in the host can limit their persistence. The target specificity of antibodies elicited in different hosts varies considerably. Whether some specificities are shared and have affected the population-level evolution of Env structure is still unclear. We examined the historical changes in amino acid sequence of the gp41 fusion peptide proximal region (FPPR), which is not exposed on the Env trimer. For three FPPR positions, the residue found in the clade B ancestor was mainly replaced by alanine. However, the changes in alanine frequency at these positions between 1979 and 2016 followed different patterns; two positions maintained a historically-constant frequency whereas the third showed a gradual increase. To understand these patterns, we introduced alanine substitutions in the FPPR of primary HIV-1 strains and examined their fitness and antigenicity relative to the clade-ancestral form. The evolutionary patterns could not be explained by effects on Env fitness. Instead, the FPPR variants with a historically-constant alanine frequency exhibited a unique open-at-the-base conformation of the trimer that exposes partially-cryptic epitopes. These Envs were modestly but significantly more sensitive to poorly-neutralizing sera from HIV-infected individuals than the clade-ancestral form. Our findings suggest that weakly-neutralizing antibodies targeting the base of the trimer are commonly elicited. Such low-level antibody pressures do not exert catastrophic effects on the emerging variants but rather determine their set-point frequency in the population and historical patterns of change.

**IMPORTANCE:** HIV-1 infection elicits antibodies that target the Env proteins of the virus. The specific targets of these antibodies vary between infected individuals. It is unclear whether some target specificities are shared between the antibody responses of different individuals. Our data suggest that antibodies against the base of the Env protein are commonly elicited during infection and are contained in sera with low neutralization efficacy. Such antibody pressures are weak. As a result, they do not completely eliminate the sensitive Env forms from the population, but rather maintain their frequency at a low level that has not increased during the past 40 years.

## INTRODUCTION

The envelope glycoproteins (Envs) of HIV-1 are primary targets in AIDS vaccine design (1,2). However, the Envs are not “stagnant” targets. Antigenic properties of these proteins have gradually changed during the AIDS pandemic among strains that circulate in the population (3-6). As a result, epitopes targeted by several broadly neutralizing antibodies (BNAbs) are found in decreasing proportions of viruses (3). Changes in Env are caused by errors that occur during reverse transcription of the viral genome, continuously introducing new variants in the host (7,8). Persistence of the variants and their establishment in the population is determined by the selective pressures applied on them, including: (i) fitness pressures, (ii) immune pressures, and (iii) bottlenecks that reduce diversity of virus forms transmitted to new hosts (9-12). The relative roles of these forces in determining the population level evolution of Env is unclear. Nevertheless, some evidence suggests that their combined effects are similar in different hosts. In a recent study by Han et al., the authors analyzed the population-level distribution of amino acids at different positions of Env (13). They observed that in distinct geographic regions, each Env position has evolved toward a specific frequency distribution of amino acids that replaced the regional ancestral form. The frequency distribution is specific for HIV-1 clade and is similar in monophyletic and paraphyletic lineages, suggesting that, at a population level, Env encounters similar selective pressures (13).

Fitness pressures play a major role in determining the nature of emerging variants that can persist in the host (14). Such pressures guide Env toward structurally stable states that effectively recognize the entry receptors and mediate fusion with target cells. Fitness pressures are likely similar in different individuals during the same stage of infection (14,15). By contrast, the target-specificity of immune pressures applied in different hosts varies considerably (16,17). Whether some targets are shared between antibody responses elicited in different hosts and guide evolution of Env structure in the population is unclear (16,18). That such common immune pressures may exist is suggested by the difference between conformational properties of Envs from lab-adapted strains and primary HIV-1 isolates. The Envs of primary isolates exhibit closed conformations that conceal immunogenic gp120 epitopes overlapping the coreceptor-binding site (CoR-BS) and CD4-BS (19-22). Lab adaptation of HIV-1 is usually associated with changes to more open forms that expose these epitopes (22,23). Such differences suggest that immune pressures reduce the frequency of forms that expose immunogenic epitopes. Whether similar pressures also affect the gp41 subunit, and their impacts on conformation of the trimer are still unknown.

To better understand the contribution of immune pressures to the population-level evolution of Env, we analyzed properties of emerging Env variants in a region of gp41 that exhibits a complex pattern of evolution. The fusion peptide proximal region (FPPR), also designated the polar region, is located C-terminal to the fusion peptide, extending from Env position 528 to 540 (24). In the unliganded form of Env (i.e., not bound to CD4), the FPPR appears to be conformationally flexible (25,26). Nevertheless, substitutions at some FPPR positions increase detachment of gp120 from virions (27,28), supporting the notion that this region contributes to structural stability of the trimer. We analyzed the historical changes in amino acid sequence of the FPPR among clade B viruses and tested the emerging variants for their fitness and antigenicity. We found that the emerging FPPR variant that is increasing in its frequency exhibits fitness levels and antigenic properties that are similar to the clade ancestral form. The FPPR variants that show a historically-constant frequency also show high fitness; however, they exhibit a unique “open-at-the-base” trimer conformation that exposes partially-cryptic gp41 epitopes. Such epitopes are targeted by Abs commonly elicited in HIV-infected individuals. Our findings suggest that the population-level changes in the FPPR can be explained by weak antibody pressures applied on the base of the trimer.

## RESULTS

### FPPR positions show distinct patterns of change during the 40 years of the AIDS pandemic

In the native state of Env, the FPPR resides at the base of the trimer, close to the C-terminal heptad repeat (CHR) region, and is minimally exposed (**Fig. 1A**). Analysis of 1,576 Envs from HIV-1 clade B showed limited variation in amino acid sequence at all FPPR positions except 535 (**Fig. 1B**). Cryo-EM reconstructions of soluble Env trimers suggest the FPPR is arranged as a helix or a helix-turn-helix adjacent to the CHR region (29-32). The CHR and FPPR appear to form a ring structure at the base of the trimer, which is supported by interactions between the two flanks of each FPPR and the CHRs of the same and adjacent protomers (**Fig. 1C** and **D**). At the N-terminus, Met530 of the FPPR resides in a clasp-like structure formed by three Trp residues from the CHR of the same protomer (26,30). At the C-terminal end of the FPPR, Arg542 of the N-terminal heptad repeat (NHR) region is located 2.8 Å from Glu647 in the CHR of the adjacent protomer. In contrast to the fixed positioning of the FPPR flanks, the mid-segment (residues 531-539) appears to be conformationally flexible (25,26). The contribution of this domain to the structure and function of Env is unknown.

**Figure 1.**
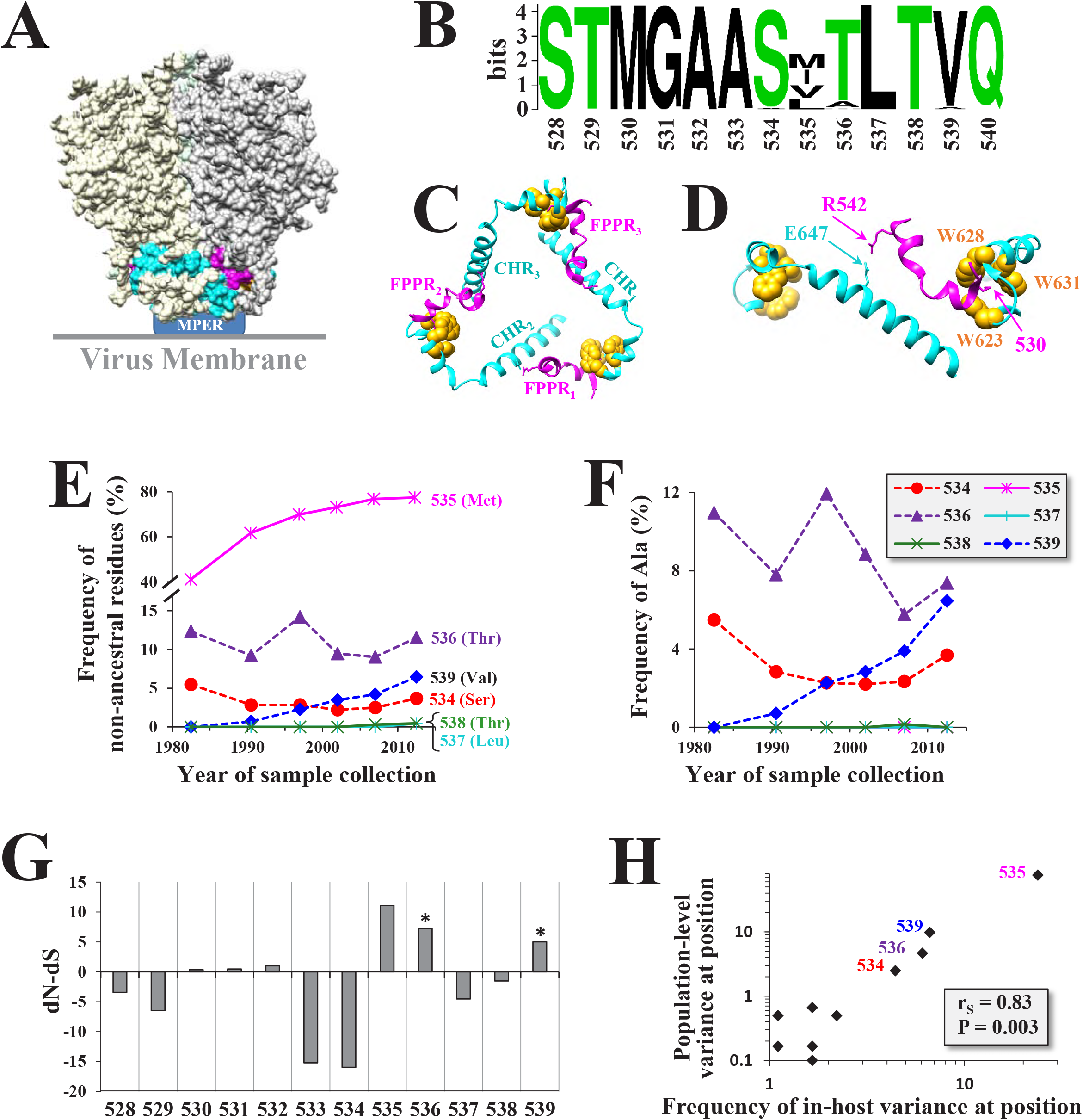
Positions in the FPPR mid-segment show distinct patterns of diversification in the population. **(A)** Cryo EM structure of the HIV-1 Env trimer (PDB ID 6CDI). The FPPR is colored in pink and the CHR in cyan. Locations of the MPER and viral membrane are shown. **(B)** Web logo of amino acid sequence distribution in the FPPR among clade B isolates circulating worldwide. Data describe 1,576 HIV-1 strains isolated from samples collected between 1979 and 2016. **(C,D)** Relationships between the FPPRs and CHR regions of the three gp41 protomers demonstrating the ring structure at the base of the trimer. The three residues of the Trp clasp are colored in yellow **(E)** Historical changes in amino acid sequence diversity at FPPR positions 534-539 among clade B isolates. Each data point describes the frequency of isolates with non-ancestral residues for the indicated 5-7-year period (as a percent of all strains from that period). The clade-ancestral residues are shown in 3-letter code. **(F)** Historical changes in frequency of Ala in clade B isolates at positions 534-539. **(G)** Positive selection in the FPPR. Estimated rates of non-synonymous (dN) and synonymous (dS) substitutions were calculated for each codon of the FPPR using 6,285 clade B sequences. The test statistic indicates the difference between dN and dS values. Asterisks indicate the P value for rejecting the null hypothesis of neutral evolution: *, P<0.05. **(H)** Correlation between in-host variance and population-level diversity of amino acid sequence at FPPR positions. A panel of 4,252 Env sequences from 181 patients were analyzed; for each patient sample 10-80 Env sequences were examined. The percent of patients that contain variation in amino acid sequence at each position is shown. This value is compared with the frequency of non-ancestral residues at the same position among clade B Envs isolated from samples collected worldwide between 2007 and 2016.

To understand the forces that act on the FPPR mid-segment, we examined the historical changes in amino acid diversity at positions 534 to 539 (**Fig. 1E**). Positions 531-533 and 540 were not included due to lack of substantial diversity. Clade B Envs isolated from samples collected worldwide between 1979 and 2016 were analyzed (13). For consecutive 5-7-year periods, we calculated at each position the frequency of emerging variants (i.e., non-ancestral residues). Position 535 showed a rapid increase in the frequency of emerging variants during the pandemic. Position 539 showed a gradual increase; the clade-ancestral Val was mainly replaced by Ala (**Fig. 1F**). At positions 534 and 536, the ancestral residues were also replaced by Ala; however, diversity and Ala frequency remained constant or decreased during the past 40 years. The emerging Ala variants at positions 534, 536 and 539 did not localize to specific lineages of the virus (see annotated phylogenetic trees in **Supplemental. Fig. S1**).

To better understand these patterns, we examined FPPR codons for evidence of positive or negative selection. Nonsynonymous (dN) and synonymous (dS) substitution rates were calculated for 6,285 HIV-1 clade B isolates. The dN-dS statistic was used as an indicator of adaptive evolution (dN-dS>0) or purifying selection (dN-dS<0). Positions 536 and 539 showed high positive dN-dS values, which were statistically significant, whereas the dN-dS value for position 534 was negative but was not significant to suggest purifying selection (**Fig. 1G**). To further investigate the selective pressures applied on FPPR positions, we examined the in-host variance in amino acid sequence at each position of this domain. We previously showed that the level of in-host variance in Env features is conserved among different hosts (3). In-host variance in amino acid sequence at each position of Env correlates well with its population-level diversity. We examined the in-host variance in amino acid sequence at each FPPR position using a panel of 4,252 Env sequences isolated from blood samples of 181 individuals (10-80 Env sequences were analyzed per sample). For each position of the FPPR, we determined the percent of patient samples that show sequence variation. This value was compared with the population-level diversity at each position, as measured by the percent of strains that did not contain the clade B ancestral residue. We observed that the frequency of in-host variance at each position correlated well with its population-level diversity (**Fig. 1G**). This finding suggests that the selective pressures applied in the host on FPPR positions determine their patterns of change in the population.

Therefore, S534, T536 and V539 in the clade B ancestor were replaced primarily by Ala. However, the three positions show different patterns of historical changes in Ala frequency in the population; 539A is gradually increasing whereas 534A and 536A have remained constant (or decreased) during the pandemic. Positions 536 and 539 show evidence of positive selection, potentially due to in-host pressures applied on these sites.

### Population-level changes in amino acid sequence of the FPPR are not fully explained by fitness of the emerging variants

To better understand the selective pressures applied on the FPPR, we examined the effects of Ala substitutions at positions 534-539 on fusion competence of Env. The changes were introduced in the Envs of two primary HIV-1 strains: (i) the Env of strain AD8, which occupies a “closed” conformation and exhibits a neutralization-resistant profile of a Tier-2 or Tier-3 virus (33-35), and (ii) the Env of strain 89.6, which is partially open and exhibits a neutralization-sensitive profile between Tiers 1 and 2 (33). Replication-defective pseudoviruses that contain the Env variants and express the luciferase protein were generated and tested for their infectivity. Substitutions to Ala at positions 536 and 539 resulted in modest but significant increases in infection relative to the wild-type (WT) forms of both strains that contain the clade B ancestral residues (**Fig. 2A**). The change at position 534 caused a modest decrease, which was not statistically significant. Other positions, which do not contain Ala in primary isolates that circulate in the population, showed varying levels of decrease in infectivity relative to the WT Envs.

**Figure 2.**
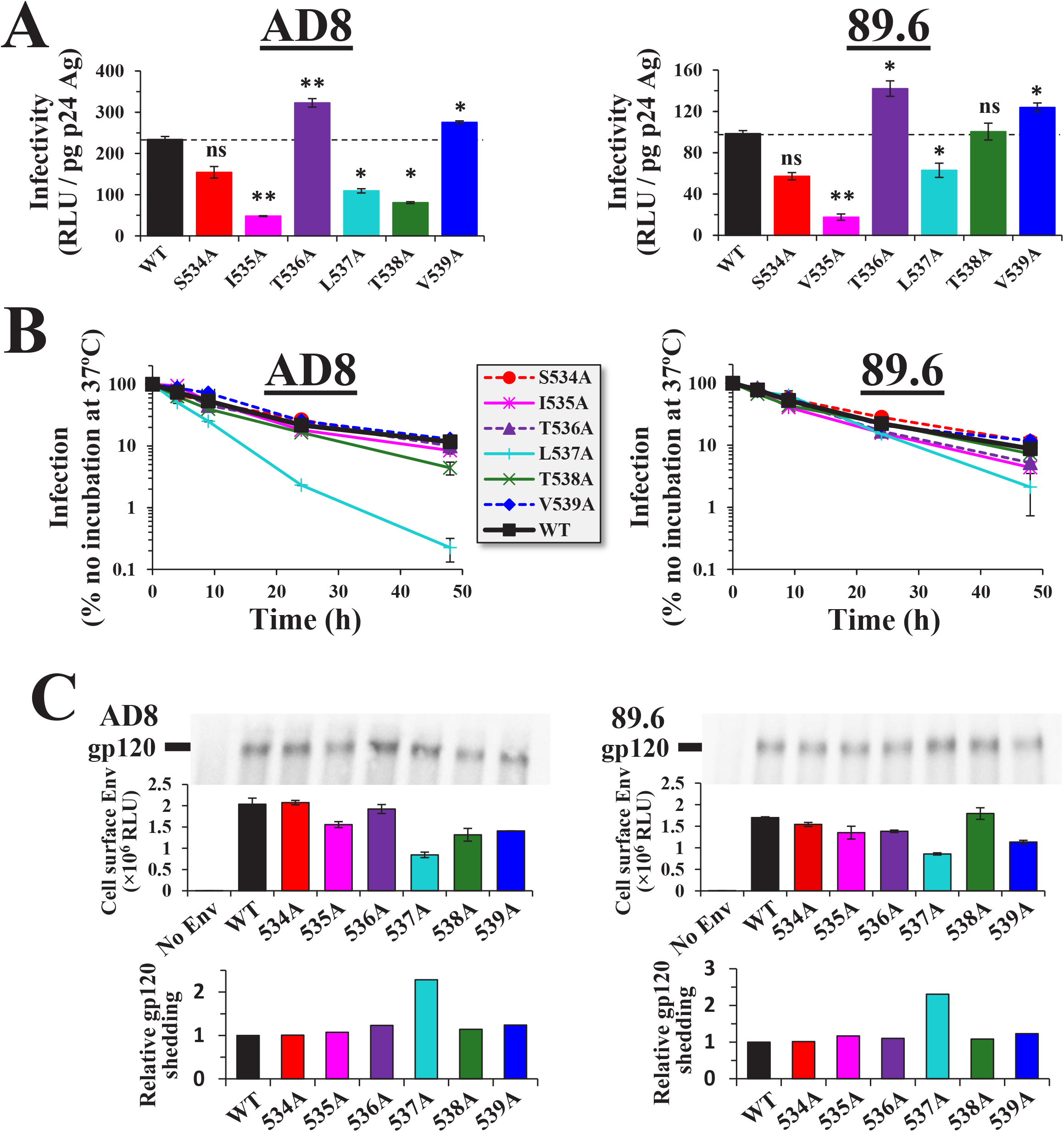
Fitness of Ala variants in the FPPR mid-segment does not fully explain their emergence patterns in the population. **(A)** Infectivity of viruses that contain Envs with Ala substitutions at positions 534-539. Replication-defective viruses that express the luciferase protein and contain the indicated variants of the AD8 and 89.6 Envs were used to infect Cf2Th-CD4^+^CCR5^+^ cells. Infectivity was measured by luciferase activity 3 d later (in relative light units, RLUs). To correct for virus particle content, infectivity is expressed as a fraction of p24 antigen in each sample. Error bars, standard error of the means (SEM). Statistical significance of the differences between infectivity of WT Env and the Ala variants was calculated using an unpaired T test that compared 5 different stocks of the viruses: *, P < 0.05; **, P < 0.005. **(B)** Stability of FPPR variants at 37°C. Frozen aliquots of luciferase-expressing viruses that contain the indicated Envs were thawed at different time points, incubated at 37°C for the indicated periods and added to the Cf2Th-CD4^+^CCR5^+^. Infectivity was measured 3 d later. Values are expressed as the percent of infection measured in samples not pre-incubated at 37°C. **(C)**Shedding of gp120 from cells that express WT and mutant Envs. HOS cells were transfected by the indicated Env variants. Three days later, the supernatant was collected and the gp120 content was immunoprecipitated using Protein A beads and mAbs 2G12, VRC03 and PGT121. Samples were analyzed by SDS-PAGE and the blot probed with goat anti-gp120 IgG. To normalize for cell-surface expression of each Env, values are expressed relative to the binding of the above mAb mix to HOS cells that express the Env variants, as measured by ELISA. The ratio between shed gp120 and cell-surface Env is expressed relative to the WT Envs (assigned a value of 1).

We also examined effects of the substitutions on Env trimer stability. For this purpose, sensitivity of the viruses to incubation at 37°C was measured. Primary strains of HIV-1 exhibit similar sensitivities to this treatment; half-lives at 37°C usually range between 7 and 10 hours (36). Ala substitutions at positions 534, 535, 536, 538 and 539 did not alter sensitivity to 37°C, whereas 537A was considerably more sensitive than the WT Envs (**Fig. 2B**). We further tested Env stability by measuring effects of the FPPR substitutions on spontaneous shedding of gp120 from Env trimers. For this purpose, HOS cells were transfected by the WT and mutant Envs. This cell type only expresses the fully cleaved form of Env on its surface (37). Shed gp120 was immunoprecipitated from the supernatant and quantified as a fraction of Env expressed on the surface of the cells. Relative to the WT Envs, only the 537A variants exhibited higher levels of gp120 shedding (**Fig. 2C**), which is consistent with stability of these variants at 37°C (**Fig. 2B**).

The above results raised the following question: If infectivity and stability of the 536A and 539A variants (and possibly 534A) are similar to or higher than the ancestral form, why is the frequency of 539A increasing in clade B whereas the frequencies of 534A and 536A are constant or decreasing? We hypothesized that the lack of an increase in frequency of 534A and 536A may reflect their sensitivity to immune selective pressures.

### Substitutions in the FPPR mid-segment induce a unique open-at-the-base conformation of the Env trimer

We examined whether the population-level changes in the FPPR can be explained by antibody (Ab) pressure applied on the emerging variants. For this purpose, we analyzed the neutralization profiles of Ala variants at positions that exhibit intermediate and high levels of infectivity: 534, 536, 537, 538 and 539. Neutralization by monoclonal Abs (mAbs) that target distinct gp41 and gp120 epitopes was compared with that of the WT Envs. We first examined sensitivity to mAbs that target the membrane-proximal external region (MPER) of gp41. MPER epitopes are partially exposed on the unliganded form of Env (38). Ala substitutions at all FPPR positions except 539 increased sensitivity to mAbs 10E8 and 2F5 (**Fig. 3A** and **B**). Similar effects of the changes were observed for strains AD8 and 89.6 (see IC_50_ values in **Table 1**). We also examined sensitivity of the variants to mAb 35O22, which targets an epitope that overlaps the gp41-gp120 interface at the base of the trimer (39,40). All FPPR variants of AD8 Env except 539A were more sensitive than the WT Env to mAb 35O22 (**Fig. 3C**). Strain 89.6 is naturally resistant to mAb 35O22. In contrast to the higher sensitivity of the FPPR variants to mAbs that target gp41, their sensitivity to mAbs that target exposed gp120 epitopes was generally similar to that of the WT Envs, including (i) mAb PGT145, which targets a quaternary epitope at the apex of the trimer (41), (ii) mAb VRC01, which targets the CD4-BS (42), and (iii) mAb PGT121, which targets an epitope that overlaps the high-mannose patch of gp120 (43) (**Fig. 3D-3F**).

**Table 1.** Sensitivity of FPPR variants to neutralization by Env-targeting mAbs.

|  |  | *2F5<sup>a</sup>* | *10E8* | *35O22* | *PGT121* | *PGT145* | *VRC01* | *17b* |
| --- | --- | --- | --- | --- | --- | --- | --- | --- |
| **89.6** | WT<sup>b</sup> | 0.41 | 0.21 | >3 | 0.02 | >2 | 2 | 22 |
|  | S534A | 0.27 | 0.1 | >3 | 0.02 | >2 | 1.9 | 3.1 |
|  | T536A | 0.06 | 0.05 | >3 | 0.02 | >2 | 0.9 | 0.4 |
|  | L537A | 0.07 | 0.04 | 0.06 | 0.02 | >2 | 1.9 | 0.8 |
|  | T538A | 0.3 | 0.1 | >3 | 0.02 | >2 | 1.9 | 19 |
|  | V539A | 0.5 | 0.21 | >3 | 0.02 | >2 | 1.7 | 18 |
| **AD8** | WT | 1.8 | 0.5 | 0.1 | 0.04 | 0.04 | 0.5 | >30 |
|  | S534A | 0.17 | 0.2 | 0.04 | 0.01 | 0.07 | 0.5 | >30 |
|  | T536A | 0.35 | 0.13 | 0.04 | 0.04 | 0.04 | 0.3 | >30 |
|  | L537A | 0.08 | 0.04 | 0.04 | 0.02 | 0.03 | 0.4 | 0.4 |
|  | T538A | 0.07 | 0.11 | 0.03 | 0.03 | 0.07 | 0.4 | >30 |
|  | V539A | 1 | 0.4 | 0.06 | 0.05 | 0.04 | 0.3 | >30 |

<sup>a</sup> IC<sub>50</sub> values are expressed in in µg/mL.
<sup>b</sup> WT, wild-type Env of HIV-1 strains 89.6 or AD8.

**Figure 3.**
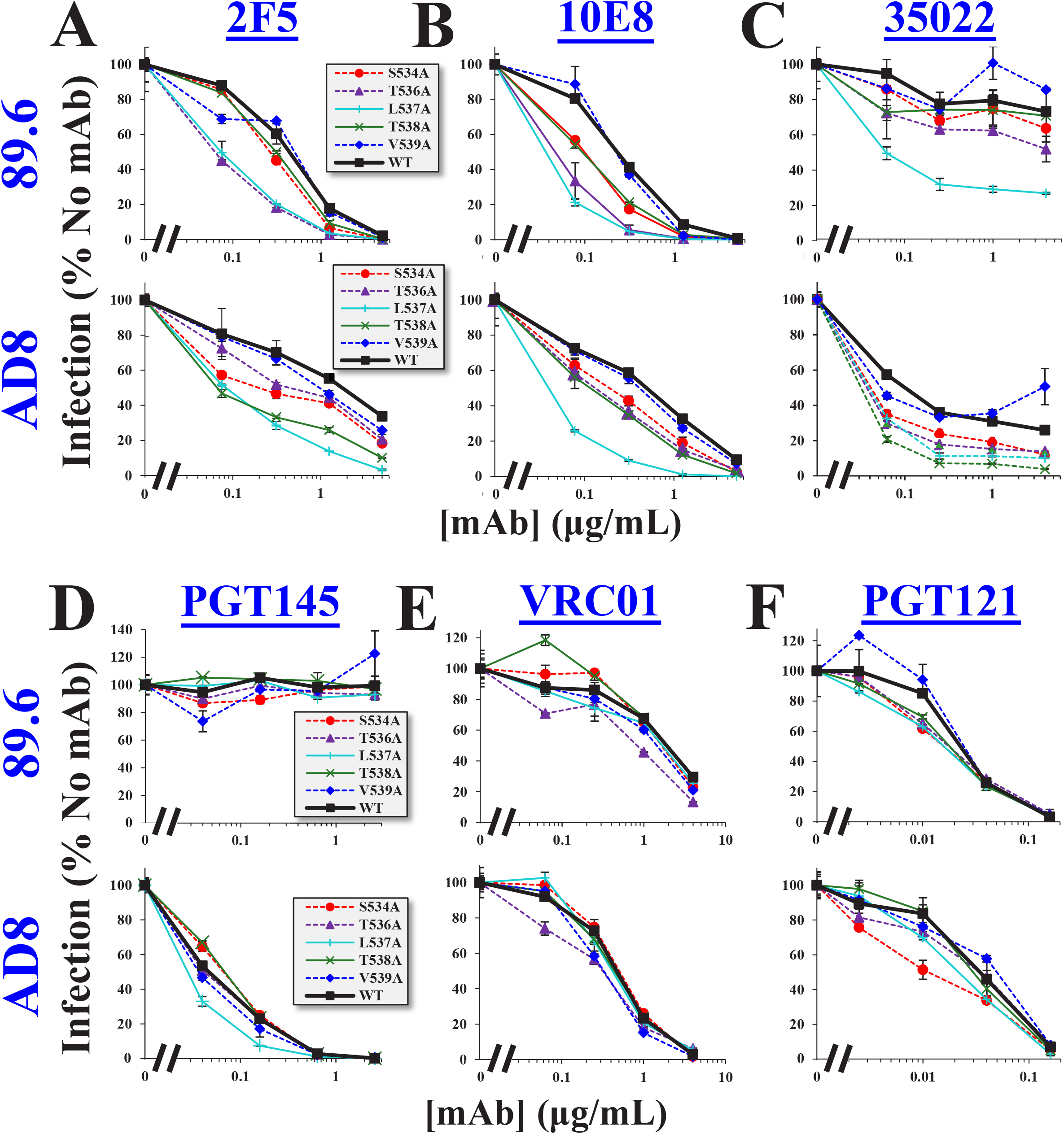
Alanine substitutions in the FPPR mid-segment induce an open-at-the-base conformation of the Env trimer. **(A-F)** Sensitivity of FPPR variants to neutralization by mAbs. Viruses that contain the Envs of strains 89.6 or AD8 with the indicated substitutions were incubated with different concentrations of the mAbs for 1 h. Samples were then added to Cf2Th-CD4^+^CCR5^+^ and residual infectivity measured 3 d later. Values describe the mean infection measured for each virus as a percent of infection measured in the absence of mAbs. Error bars, SEM.

We also examined sensitivity of the Envs to mAb 17b, which targets a cryptic epitope that overlaps the CoR-BS (44). For HIV-1 AD8, only the 537A variant was neutralized by this mAb (**Fig. 4A**). WT 89.6 Env is partially open and, similar to many Tier-1 strains, is modestly sensitive to CoR-BS-targeting Abs (33) (**Fig. 4A**). The 534A, 536A and 537A changes in 89.6 Env increased sensitivity to mAb 17b, suggesting that they further enhanced exposure of the CoR-BS. Since Envs of most primary HIV-1 strains occupy a more closed conformation than 89.6, we sought to determine if introducing these substitutions in a closed form of this Env would also cause exposure of the CoR-BS. For this purpose, we used a variant of 89.6 that contains two stabilizing changes in gp120: (i) Arg to Glu at position 305 of the V3 loop, and (ii) Met to Ile at position 225 of the gp120 inner domain (45,46). These changes were identified after serial passage in rhesus macaques of a chimeric simian-human immunodeficiency virus that contains the 89.6 Env (47). Analysis of the antigenic profile of 89.6(M225I,R305E) Env showed it occupies a more closed conformation than the parental 89.6 strain (33). We introduced the FPPR changes 534A and 536A in the 89.6(M225I,R305E) Env and tested sensitivity to different mAbs. As expected, 89.6(M225I,R305E) was more resistant than WT 89.6 to mAb 17b (IC_50_ values were >60 μg/mL and 25 μg/mL, respectively; compare **Fig. 4A** and **4B**). Importantly, both 534A and 536A changes increased sensitivity of 89.6(M225I,R305E) to MPER mAb 2F5 but did not alter sensitivity to mAb 17b.

**Figure 4.**
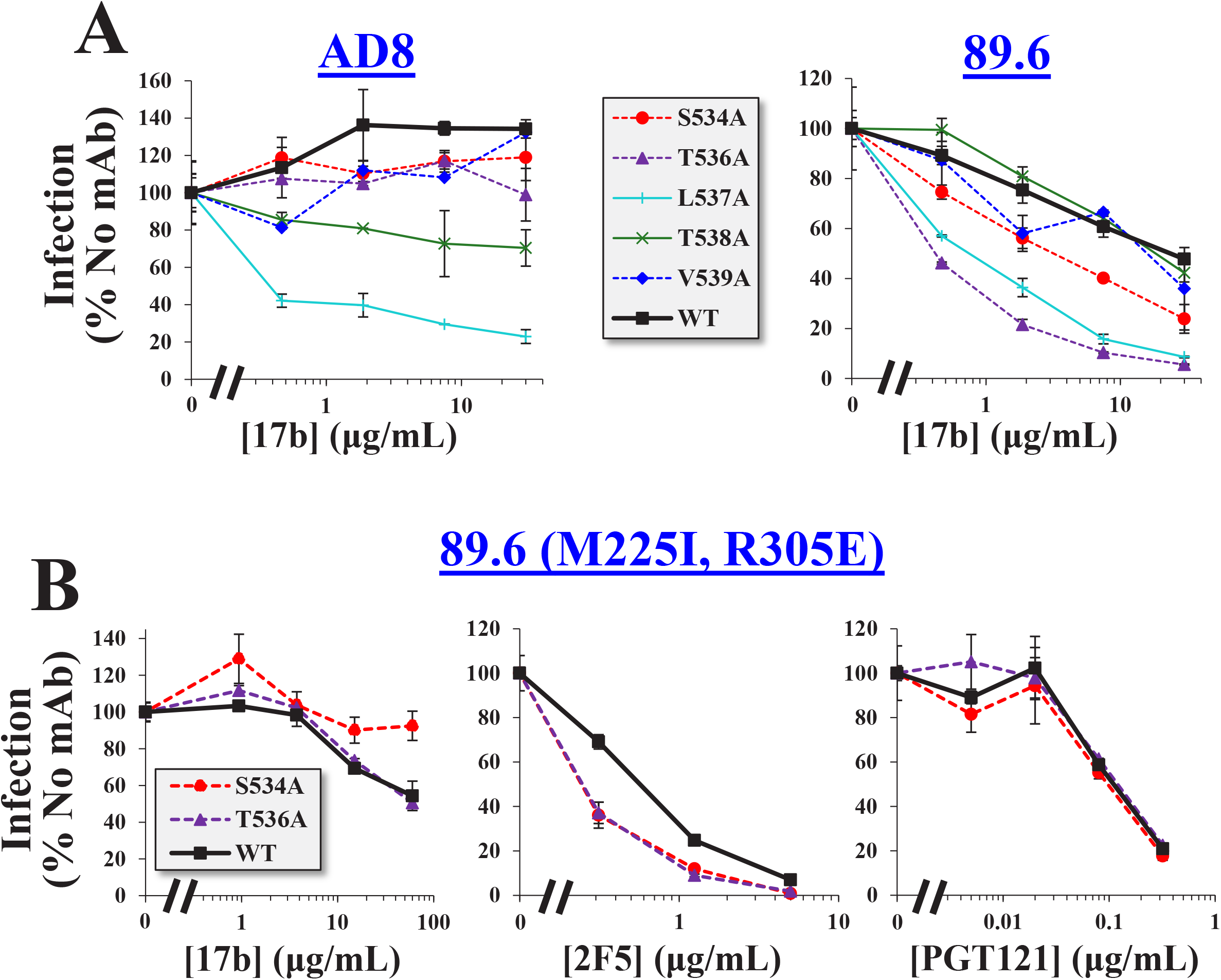
Ala substitutions in the FPPR of a Tier-3-like variant of 89.6 Env enhance exposure of the MPER but not the CoR-BS. **(A)** Sensitivity of viruses containing the indicated variants of strains 89.6 or AD8 to mAb 17b. **(B)** Effects of Ala substitutions in the FPPR on sensitivity of Env 89.6(M225I,R305E) to mAbs 17b, 2F5 and PGT121. Error bars, SEM.

The above data suggest that the FPPR regulates exposure of epitopes at the trimer base. In primary Tier-2/3-like viruses (i.e., AD8 and 89.6(M225I,R305E)), Ala substitutions at FPPR positions 534 and 536 induce isolated exposure of epitopes in the MPER and gp120-gp41 interface. Ala substitutions at other FPPR positions (or in partially-open forms, such as 89.6 Env) can induce more drastic changes in trimer arrangement and enhance exposure of epitopes that overlap the CoR-BS.

### FPPR substitutions that induce an open-at-the-base conformation sensitize HIV-1 to antibodies commonly elicited during HIV-1 infection

The emerging variants 534A and 536A show a historically-constant frequency in the population and exhibit an open-at-the-base conformation of the trimer. By contrast, the 539A variant that is increasing in its frequency exhibits a more closed conformation. We asked whether common Ab pressures applied on the open-at-the-base forms may account for their lack of increase in the population. To address this question, we tested sensitivity of the above variants to serum samples from HIV-1-infected individuals. We first examined sensitivity of the variants to three pools of serum, each composed of samples from 10-16 HIV-1-infected individuals. As expected, WT AD8 was more resistant to the patient sera than 89.6 (compare top and bottom panels in **Fig. 5**). Considerable variation was observed in effects of the Ala substitutions on sensitivity to the sera. Other than AD8(L537A), which was more sensitive than WT AD8 to all three pools of serum, other variants did not show consistently higher sensitivity.

**Figure 5.**
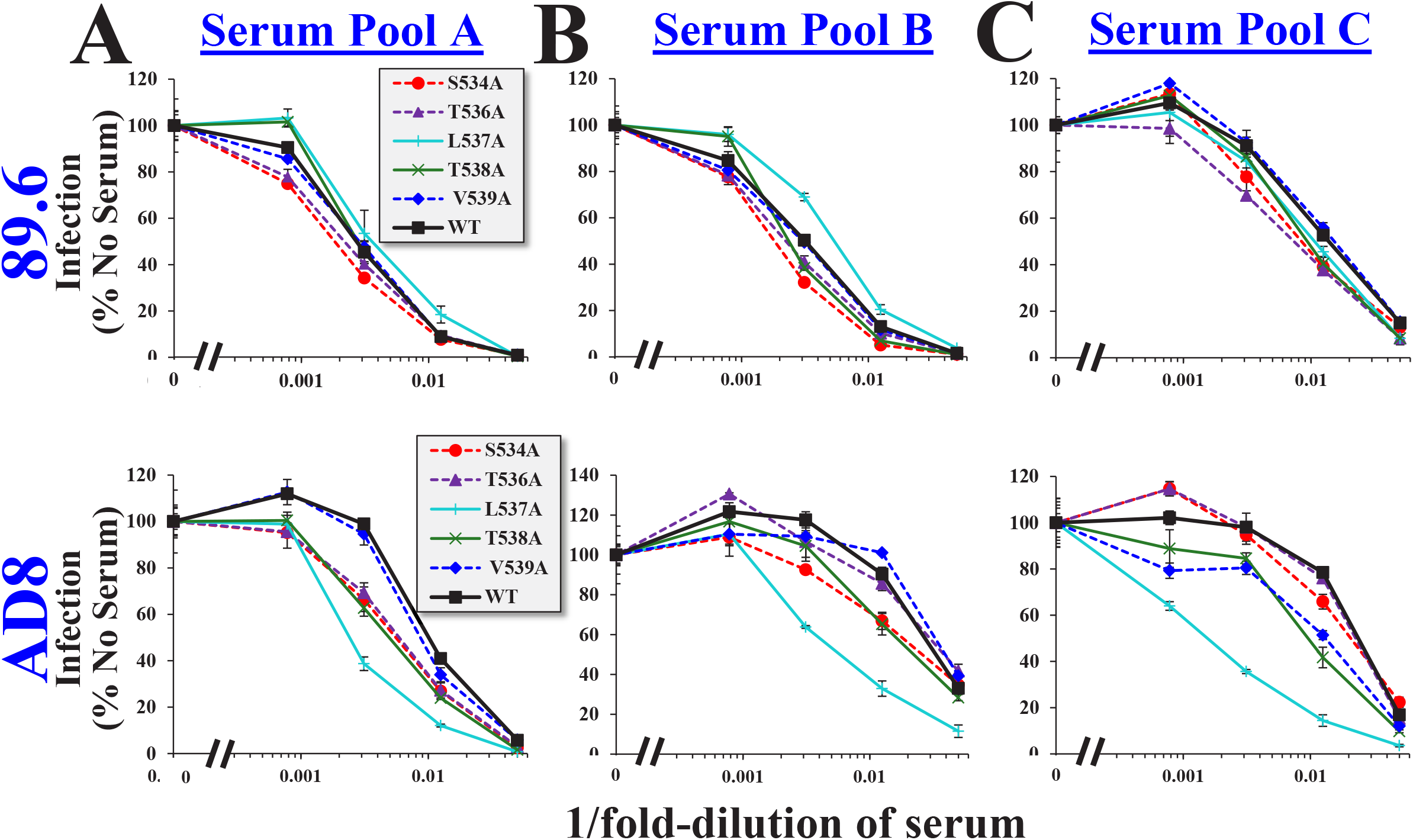
Sensitivity of FPPR variants to three pools of serum from HIV-infected individuals. Viruses that contain the indicated variants of Envs 89.6 or AD8 were incubated with different dilutions of pooled serum (each composed of samples from 10-16 individuals) and added to Cf2Th-CD4^+^CCR5^+^ cells. Infectivity was measured 3 d later and is expressed as a percent of infection measured for viruses not pre-incubated with serum. Error bars, SEM.

Since single samples that contain highly potent Abs can dominate the neutralization profiles of serum pools, we tested the effects of the substitutions on sensitivity to individual serum samples, each collected from a different donor. Samples from 14 patients that exhibit low potency (IC90 values at a dilution of 1:10 or lower, as measured for virus containing WT AD8 Env) were tested for their neutralization of the WT Envs, as well as variants 534A, 536A and 539A (**Fig. 6**). In both AD8 and 89.6 strains, the 534A and 536A variants were modestly but significantly more sensitive to the sera than the WT Envs (see results of paired T tests in **Fig. 6A** and **6B**). By contrast, sensitivity of the 539A variant did not differ from that of the WT Env. These data suggest that Abs targeting the open conformation of the 534A and 536A variants are commonly elicited in HIV-1-infected individuals and contained in samples with low neutralization efficacy.

**Figure 6.**
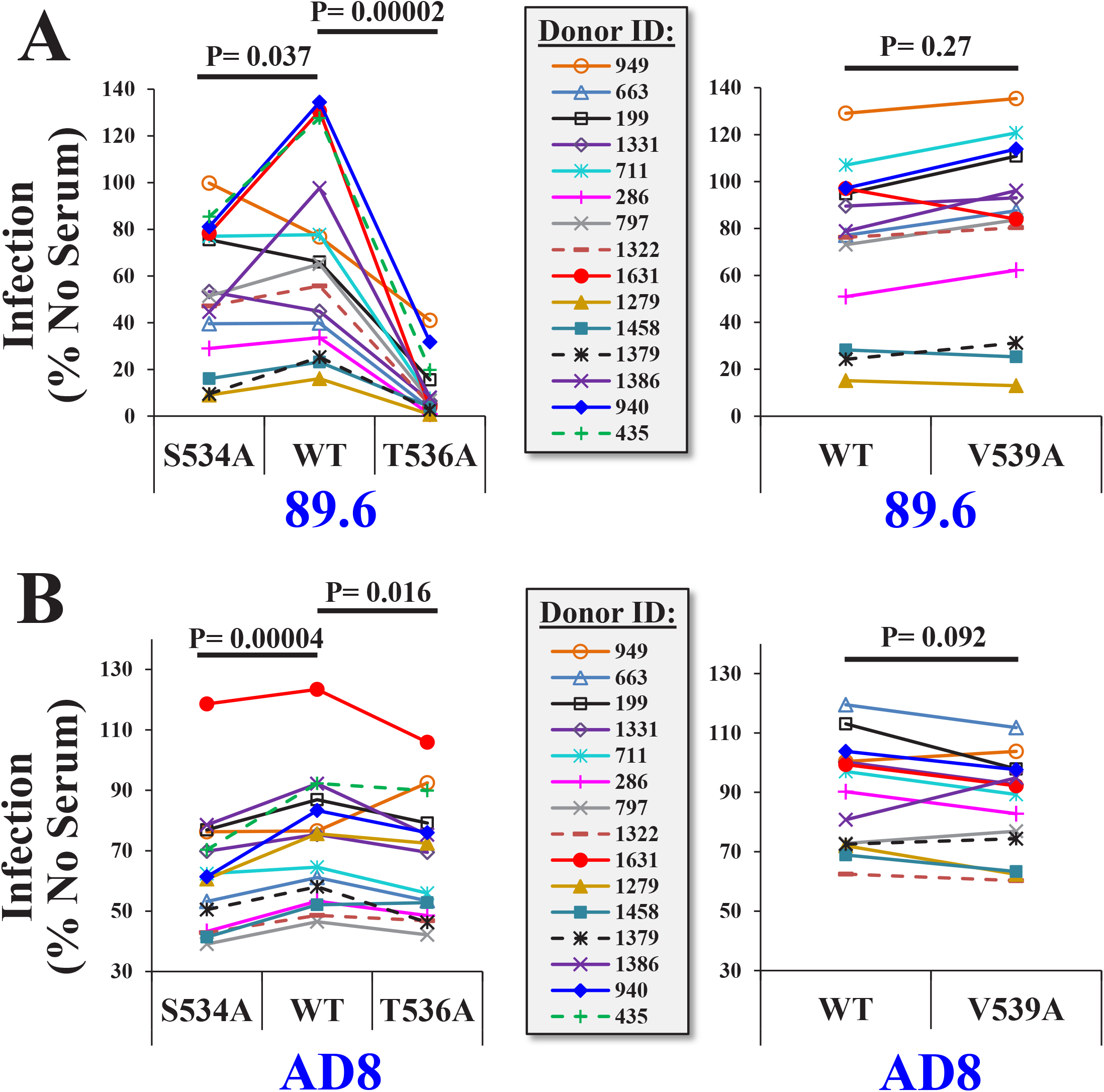
Ala substitutions in the FPPR enhance sensitivity to weakly-neutralizing serum from HIV-infected individuals. Samples from 14 individuals that exhibit low neutralization efficacy (IC90 value at a 1:10 dilution or lower for HIV-1AD8) were incubated with the WT, S534A, T536A or V539A variants of Envs AD8 and 89.6. All serum samples were used at a final dilution of 1:10. The virus-serum mix was incubated for 1 h at 37°C and added to Cf2Th-CD4^+^CCR5^+^ cells to measure residual infectivity. P values describe the results of a paired T test that compares infectivity of the WT and mutant Envs in the presence of each serum.

## DISCUSSION

Multiple sites in the Env proteins show considerable diversity in amino acid sequence among primary HIV-1 strains whereas others are highly conserved. In many cases, conservation results from fitness pressures applied on the sites, allowing only specific Env variants to persist due to their functionality or structural stability (48,49). Over the course of multiple replication cycles, even small reductions in fusion competence (as measured *in vitro*) can result in rapid loss of the variant from the host and thus limited representation in the population. However, in some cases, Env variants exhibit *in vitro* fitness levels that are similar to (or higher than) the ancestral form but are still not found among circulating strains. For example, several lab-adapted strains of HIV-1 exhibit higher fusion competence than their parental primary isolates, with lower or no requirement for CD4 to mediate fusion (50,51). Ability to surmount the necessity for CD4 would potentially allow more efficient replication in the host. Nevertheless, CD4-independent primary strains are rarely encountered; their absence is explained by the altered conformations of these Envs, which expose epitopes that overlap the CoR-BS (35,52,53). Abs that target the CoR-BS are commonly elicited in infected individuals (54) and likely apply strong selective pressures that account for the absence of CD4-independent strains in the population despite their potential fitness advantage. Our data suggest that a similar type of Ab selection pressure is applied on epitopes at the base of the trimer. However, in contrast to the CD4-independent strains, such effects do not cause catastrophic population size reductions; strains with an open-at-the-base conformation (e.g., 534A or 536A) are indeed isolated from infected individuals. Instead, their immune selection is incomplete, which results in a lack of increase in their frequency during the past four decades. The balance between the rate of appearance of such variants in the host, their fitness and sensitivity to selective pressures determines their “set-point frequency” in the population and historical patterns of change.

Structural integrity of the Env trimer is maintained by multiple interactions within and between subunits. Such sites control the overall architecture of the trimer and keep it in a functional closed form that does not expose immunogenic epitopes. Disruption of the interactions by mutations, chemical or physical treatments, or specific ligands can cause conformational changes to functional non-native forms. A common feature of these perturbed forms is exposure of epitopes that overlap the CoR-BS (33,35,55). This common path reflects the natural propensity of the trimer to “spring” into an open form that can bind the CoR. During the course of their evolution, primate lentiviruses likely gained their dependence upon CD4, a feature that limits exposure of the conserved immunogenic CoR-BS. Similar to the CoR-BS, the MPER is highly conserved (56) and is required for fusion (57-59). The MPER is thus maintained less accessible to targeting Abs (60,61); those BNAbs that target this domain generally have lower potencies than BNAbs that target more exposed epitopes on gp120. Here we show that exposure of MPER epitopes is controlled by the FPPR. Modestly-perturbing substitutions in the FPPR cause isolated exposure of these epitopes whereas more perturbing changes (e.g., L537A) induce “fully-open” trimer forms that also expose the CoR-BS (**Fig. 7**). Interestingly, a similar phenotype is caused by disruption of the anchoring interaction between gp41 and membrane cholesterol. Low concentrations of cholesterol-depleting agents increase exposure the MPER whereas higher concentrations also increase exposure of the CoR-BS (62). We expect that the changes in the FPPR induce similar release of structural constraints that maintain the trimer in a closed conformation. Based on proximity of the FPPR to the CHR, is it likely that the FPPR substitutions affect the interaction between the two domains (28). The nature of the association between the MPER and other components of gp41, which can explain the structural basis for our observations, remains to be clarified.

**Figure 7.**
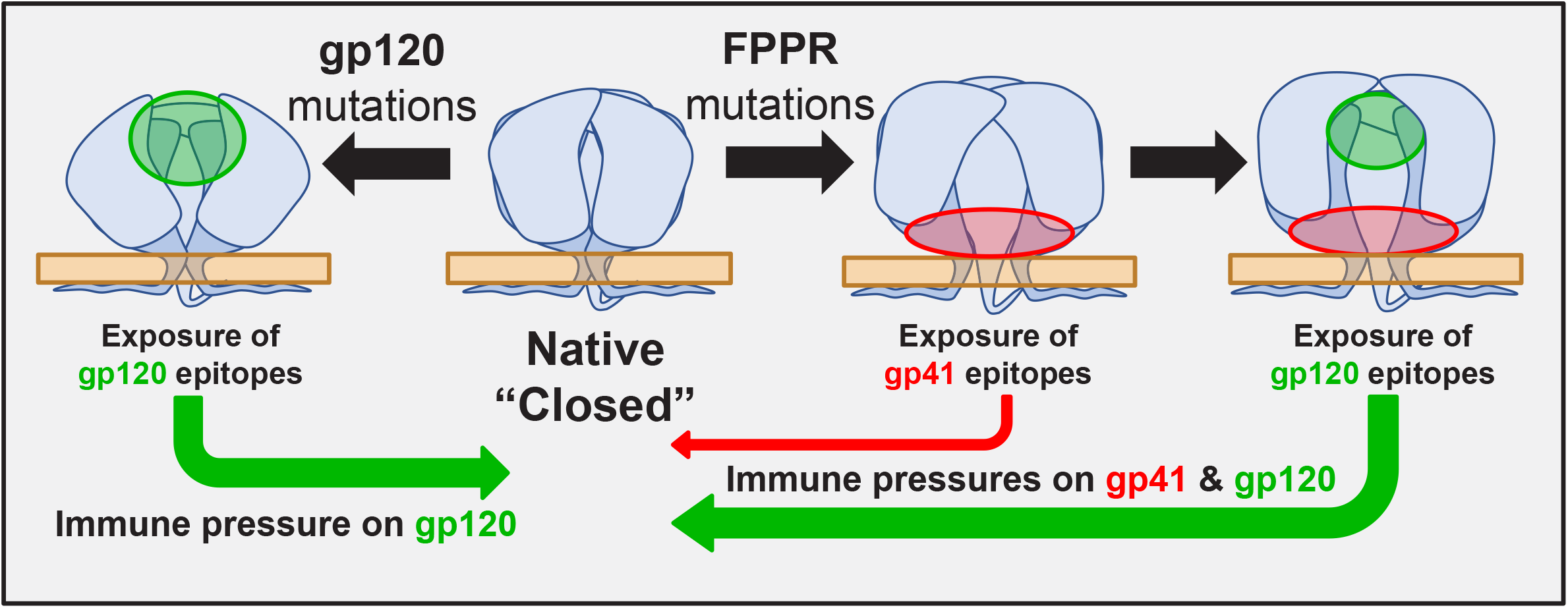
Model of the effects of immune selective pressures on the HIV-1 Env trimer. The Env trimer occupies a closed conformation with limited exposure of immunogenic epitopes in gp120 and gp41. Substitutions at different sites in the V1, V2 and V3 variable loops of Env release structural constraints and increase exposure of cryptic gp120 epitopes that overlap the CoR-BS. Strains with such open forms of gp120 are functional but are less likely to persist in the host due to their high sensitivity to neutralization by commonly-elicited Abs. Similarly, mutations at some positions in the FPPR do not reduce fitness but rather induce an open-at-the-base conformation that exposes epitopes in the gp41 MPER and gp41-gp120 interface. Weak immune pressures on these sites by commonly elicited Abs reduces the frequency of such FPPR variants in the population. Other mutations in the FPPR (e.g., at position 537) enhance, in addition, exposure of epitopes that overlap the CoR-BS; such variants are effectively eliminated and poorly represented in the population.

In different populations worldwide infected by virus from the same HIV-1 clade, each Env position exhibits a similar frequency distribution of amino acids (13). In many cases, founder effects of substitutions in monophyletic lineages are reduced by evolution of the position toward the clade-specific frequency distribution. That evolution is directed toward distributions rather than single amino acids suggests that a fine balance exists between the inherent propensity of the virus for diversification and the selective pressures that oppose it. In such cases, no single factor dominates the pattern of circulating variants. For example, the emerging variant T536A shows modestly but significantly higher fitness than the clade ancestral form but also higher sensitivity to immune pressures normally elicited in the host. Such factors are balanced and likely explain the constant frequency of this variant in the population. By contrast, other FPPR positions suggest catastrophic population reduction effects. For example, position 538 contains the same nucleotide sequence in the clade B ancestor as position 536; however, 538A variants are not found among circulating strains. In this study we show that *in vitro* analyses of fitness and potential Ab pressures can explain the distribution of emerging variants in the population and their historical patterns of change.

Broadly-acting and potently-neutralizing Abs are rarely elicited in patients. A better understanding of the lower-efficacy neutralizing responses that are more frequently elicited may provide insights that will allow us to achieve this goal. Our findings suggest that poorly neutralizing sera contain weak Ab pressures that target the trimer base, including the MPER. The high conservation and contiguous nature of the MPER render it an attractive target for immunogen design. However, previous attempts to elicit MPER-targeting Abs have yielded mixed results (63,64). The low neutralization efficacy of the elicited Abs is likely not caused by low immunogenicity of the MPER peptides used, but rather by the low exposure of the epitopes on the membrane-bound form of the Env trimer. A second limitation of MPER Abs is their propensity to associate non-specifically with the plasma membrane, resulting in autoreactivity and polyreactivity (65,66). Nevertheless, some MPER Abs have shown high levels of specificity, indicating that polyreactivity is not an inherent property of these Abs (67). That such Abs are commonly elicited in the host, as our data suggest, may indicate that additional focus on the exposed segment of this domain (68), may further increase the neutralizing efficacy of the response. In the context of an immunogen composed of the complete ectodomain of the Env trimer, exposure of such epitopes can be controlled by the FPPR.

## MATERIALS AND METHODS

### HIV-1 Env sequence analysis

To examine historical changes in the FPPR, HIV-1 *env* sequences were obtained from the Los Alamos National Lab (LANL) database using the sequence search interface (https://www.hiv.lanl.gov) and from the NCBI database (https://www.ncbi.nlm.nih.gov). Sequences tagged as nonfunctional Envs were removed, as were sequences with nucleotide ambiguities or large deletions in conserved regions. A single *env* from each patient and a single sequence from known transmission pairs were used. In addition, to avoid related sequences, we applied a minimal cutoff of 0.03 nucleotide substitutions per site for sequence selection. Nucleotide sequences were aligned using a Hidden Markov model with the HMMER3 software (69). Sequences were then translated, and positions numbered according to the standard HXBc2 numbering system of the Env protein (24). To analyze the historical changes in amino acid frequency at Env positions, sequences were grouped according to the year of sample collection and the frequency of each residue was calculated as a percent of all isolates from the same time period.

To examine the selective pressures applied on FPPR positions, we downloaded from the NCBI database the nucleotide sequence of the FPPR mid-segment (nucleotide positions 7806 to 7841 of the HIV-1 genome, corresponding to amino acid positions 528-539 of Env). A panel of 6,285 clade B *env* sequences from samples collected between 1979 and 2018 were used to estimate the number of inferred synonymous and nonsynonymous substitutions. Analyses were performed using the MEGA7 platform (70). The ancestral state of this segment was reconstructed using a Muse-Gaut model of codon substitution (71) and Felsenstein 1981 model of nucleotide substitution (72). To estimate maximum likelihood values, a tree topology was computed. To detect codons under selection, we calculated the number of synonymous substitutions per site (dS), and the number of nonsynonymous substitutions per site (dN). Maximum Likelihood computations of dN and dS were conducted using HyPhy (73). The dN-dS statistic was applied to detect codons that have undergone positive selection (dN-dS>0) or negative selection (dN-dS>0). P values for rejecting the null hypothesis of neutral evolution were calculated (74,75), whereby P values less the 0.05 were considered significant.

To calculate within-host diversity at FPPR positions, we used nucleotide sequences of 4,252 clade B Envs from 181 different HIV-infected individuals. For each individual, 10 to 80 Env sequences were analyzed. Sequences were aligned using a Hidden Markov model with the HMMER3 software (69). For each patient sample, we determined the presence or absence of in-host amino acid variation at each position of the FPPR. The percent of patient samples that contain non-zero variance at each position was determined.

### Antibodies and cells

The following mAbs were obtained through the NIH AIDS Reagent Program, Division of AIDS, NIAID, NIH. The mAb 35O22 that targets the gp120-gp41 interface was contributed by Jinghe Huang and Mark Connors (39). James Robinson provided mAb 17b, which recognizes an epitope that overlaps the CoR-BS (44). Hermann Katinger provided mAb 2F5 that recognizes the gp41 MPER (76). The MPER-targeting mAb 10E8 was contributed by Mark Connors (77). John Mascola provided the CD4-BS mAb VRC01 (78). The International AIDS Vaccine Initiative (IAVI) Neutralizing Antibody Consortium provided mAb PGT121, which recognizes a glycan-dependent gp120 epitope (43), and mAb PGT145, which recognizes a quaternary epitope at the trimer apex (43). Serum samples were obtained from HIV-1 chronically-infected adult subjects (more than 6 months since seroconversion) who gave informed consent under clinical protocols approved by the human use review boards at the University of Washington at Seattle Center for AIDS Research (CFAR) and the University of Iowa (IRB numbers 8807313, 200010008, and 2010101730). All samples were heat inactivated at 55°C for 30 min before use. Neutralization tests were conducted with individual sera and with 3 pools of serum, each composed of samples from 10-16 patients.

Human embryonic kidney 293T cells were obtained from the American Type Culture Collection (ATCC) and cultured in Dulbecco’s Modified Eagle Medium (DMEM) supplemented with 10 μg/mL penicillin/streptomycin and 10% fetal calf serum (DMEM/FCS). Canis familiaris thymus normal (Cf2Th) cells (kindly provided by Joseph Sodroski) were used to measure infection. These cells stably express human CD4 and CCR5 (Cf2Th-CD4^+^CCR5^+^) and were cultured in DMEM/FCS supplemented with 200 μg/ml hygromycin and 400 μg/ml G418.

### Envelope glycoprotein constructs

The full-length Envs of HIV-1 strains 89.6 (accession number U39362) and AD8 (accession number AF004394) were expressed from the pSVIIIenv plasmid, as previously described (33). Amino acid numbering of Env positions is based on the HXBc2 reference system (24). Mutations were introduced into the pSVIIIenv vector by site-directed mutagenesis using the PrimeStar Max polymerase (Takara), followed by DpnI digestion, and transformation of Stellar competent cells. The *env* genes of all variants generated were sequenced to verify that unwanted mutations were not introduced during this process.

### Preparation of recombinant luciferase-expressing HIV-1

Single-round, recombinant HIV-1 that expresses the luciferase gene was generated by transfection of 293T cells using JetPrime transfection reagent (Polyplus). Cells were seeded in 6-well plates (8.5 × 10^5^ cells per well) and transfected the next day with 0.4 μg of the HIV-1 packaging construct pCMVΔP1ΔenvpA, 1.2 μg of the firefly luciferase-expressing construct pHIvec2.luc, 0.4 μg of plasmid expressing HIV-1 Env, and 0.2 μg of plasmid expressing HIV-1 Rev. The transfection medium was replaced the next day and virus-containing supernatant was collected 24 h later. Supernatants were cleared of cell debris by centrifugation at 700 × *g* and filtered through 0.45-μm pore-sized membranes. Samples were snap frozen on dry ice immersed in ethanol for 15 min and stored at −80°C until use.

### Infection by luciferase-expressing HIV-1 and antibody neutralization assays

Cf2Th-CD4^+^CCR5^+^ cells were seeded in 96-well luminometer-compatible plates at a density of 7.5 × 10^3^ cells per well and infected the next day. For neutralization assays, virus preparations were incubated at 37°C in the absence or presence of mAbs or serum for 1 h. Samples were then added to Cf2Th-CD4^+^CCR5^+^ cells and incubated for 3 d to allow infection. To measure infection, the medium was removed, cells were lysed with 35 μl passive lysis buffer (Promega) and subjected to three freeze-thaw cycles. To measure luciferase activity, 100 μl of luciferin buffer (15 mM MgSO_4_, 15 mM KPO_4_ [pH 7.6], 1 mM ATP, and 1 mM dithiothreitol) and 50 μl of 1 mM D-luciferin potassium salt (Syd Labs, MA) were added to each sample. Luminescence was recorded using a Synergy H1 microplate reader (BioTek Instruments).

To measure virus sensitivity to inactivation at 37°C, recombinant viruses were generated as described above, diluted and divided into aliquots (one sample for each prospective time point). All samples were then snap-frozen on dry ice immersed in ethanol for 15 min and stored at −80°C. At different time points, the samples were thawed in a 37°C water bath for 2 min and then further incubated at 37°C for different time periods. All samples were subsequently added to Cf2Th-CD4^+^CCR5^+^ cells, and infectivity was measured 3 d later by luciferase activity.

### p24 antigen assay to quantify virus particle content

To normalize measured infectivity values by virus particle content in the samples, we quantified p24 antigen levels. For this purpose, human anti-HIV-1 p24 antigen mAb was incubated at 1 μg/mL in luminometer-compatible 96-well protein-binding plates overnight. The wells were then washed with blocking buffer, composed of 150 mM NaCl, 3 mM Tris pH8, 1.8 mM CaCl2, 1 mM MgCl_2_ and 2% bovine serum albumin (BSA). To lyse virions, samples were supplemented with 0.5% Triton X and incubated at 100°C for 5 min. Samples were then added to the 96-well plate and incubated for 2 h at room temperature. Wells were then washed 4 times with blocking buffer and a rabbit anti-p24 antigen mAb suspended in blocking buffer was added. Binding of the latter was detected by a horseradish peroxidase (HRP) conjugated goat anti-rabbit polyclonal Ab and measured by luminescence using SuperSignal West Pico enhanced chemiluminescence reagents and a Synergy H1 microplate reader.

### Shedding of gp120 from Env-expressing cells

To quantify shedding of gp120, HOS cells were seeded in 6-well plates (2.75 × 10^5^ cells per well) and transfected the next day with plasmids expressing Env, Tat and Rev using 0.4, 0.33 and 0.2 μg of each plasmid per well, respectively, and JetPrime transfection reagent. The next day samples were washed twice with Tris-Saline buffer (140 mM NaCl, 1.8 mM CaCl2, 1 mM MgCl2 and 25 mM Tris, pH 7.5) and then FreeStyle 293 Expression Medium (Gibco) was added. Two days later, the supernatant was collected, cleared of cell debris by centrifugation at 700 × *g* and filtered through 0.45-μm pore-sized membranes. Envs were immunoprecipitated using Protein A beads and a combination of mAbs 2G12, VRC01 and PGT121 (all at 1 μg/ml). Samples were then analyzed by SDS-PAGE, transferred to PVDF membranes, probed with goat-anti-gp120 and detected by an HRP-conjugated rabbit anti-goat Ab. The amount of gp120 shed into the medium was normalized for the relative expression level of each Env. For this purpose, HOS cells were seeded in 96-well plates (1.4 × 10^4^ cells per well) and transfected the next day with plasmids expressing Env, Tat and Rev, using 60, 11 and 6 ng of each plasmid per well, respectively and JetPrime reagent. Three days later, cells were washed in Tris-Saline buffer containing 2% BSA (TS/BSA) and incubated with the above combination of mAbs (all at 1 μg/ml) in TS/BSA for 1 h at room temperature. Cells were then washed twice with TS/BSA and once with TS. Binding was detected using HRP-conjugated goat-anti-human IgG and measured by luminescence using SuperSignal West Pico reagents and a Synergy H1 microplate reader. Intensity of the gp120 band from the supernatant (quantified by densitometry) was compared with the expression level of each Env, and the ratio between the two values was used to quantify gp120 shedding.

## Acknowledgments

This work was partly supported by amfAR grant 110028-67-RGRL to HH and by NIH grants AI141495 and AI150343 to LW. The authors have no conflicting financial interests.

## SUPPLEMENTAL INFORMATION

**Figure S1.**
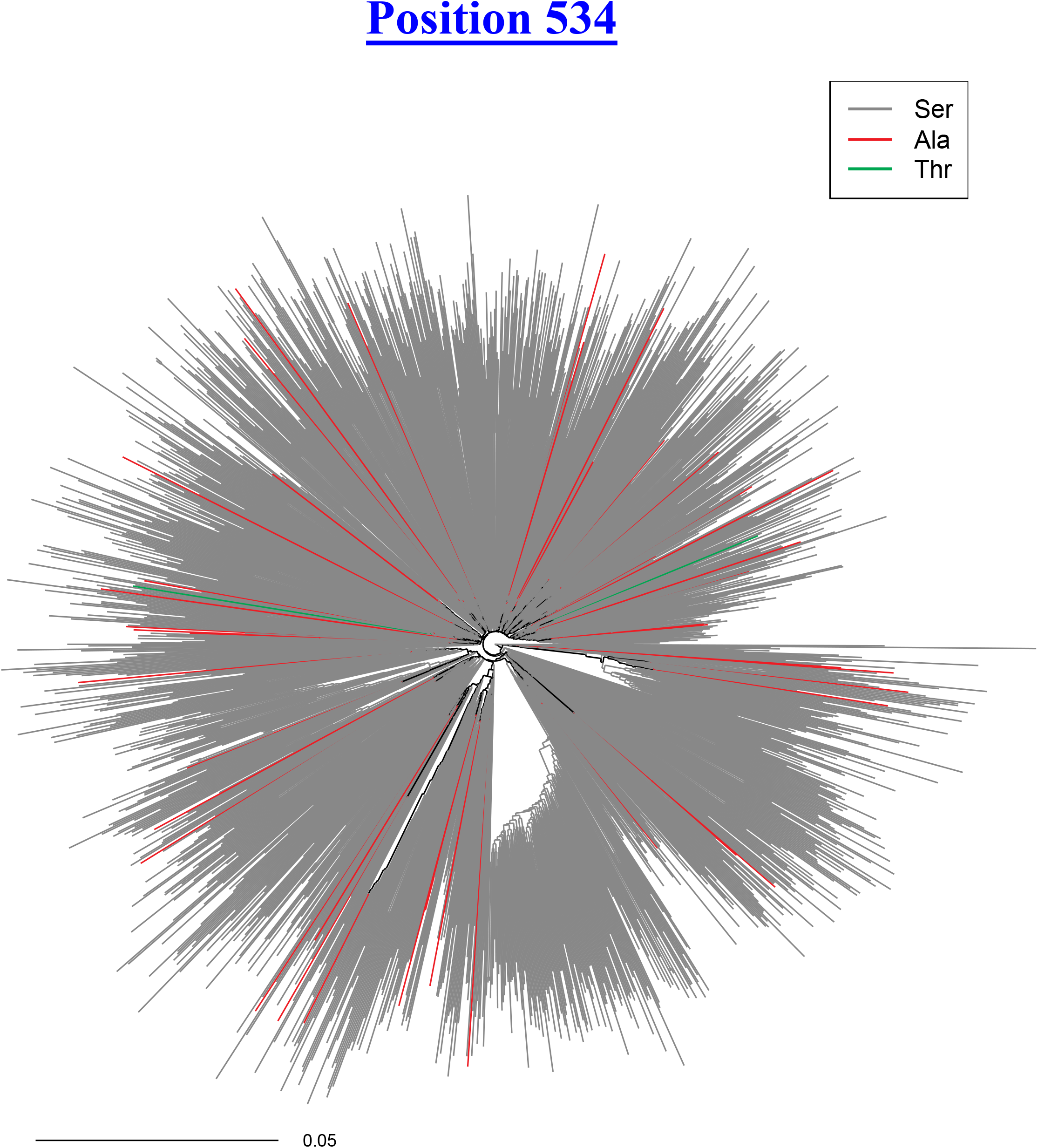
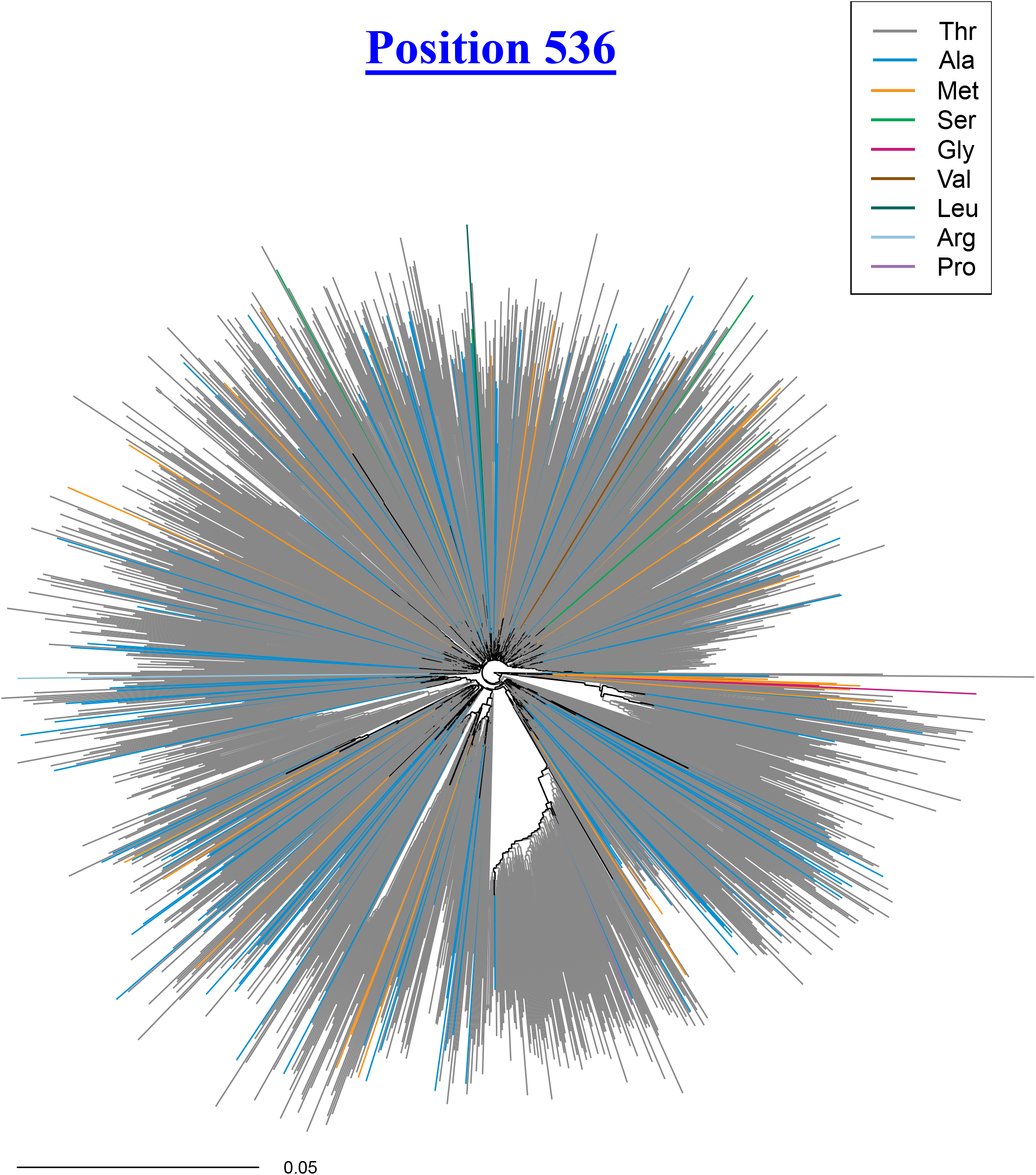
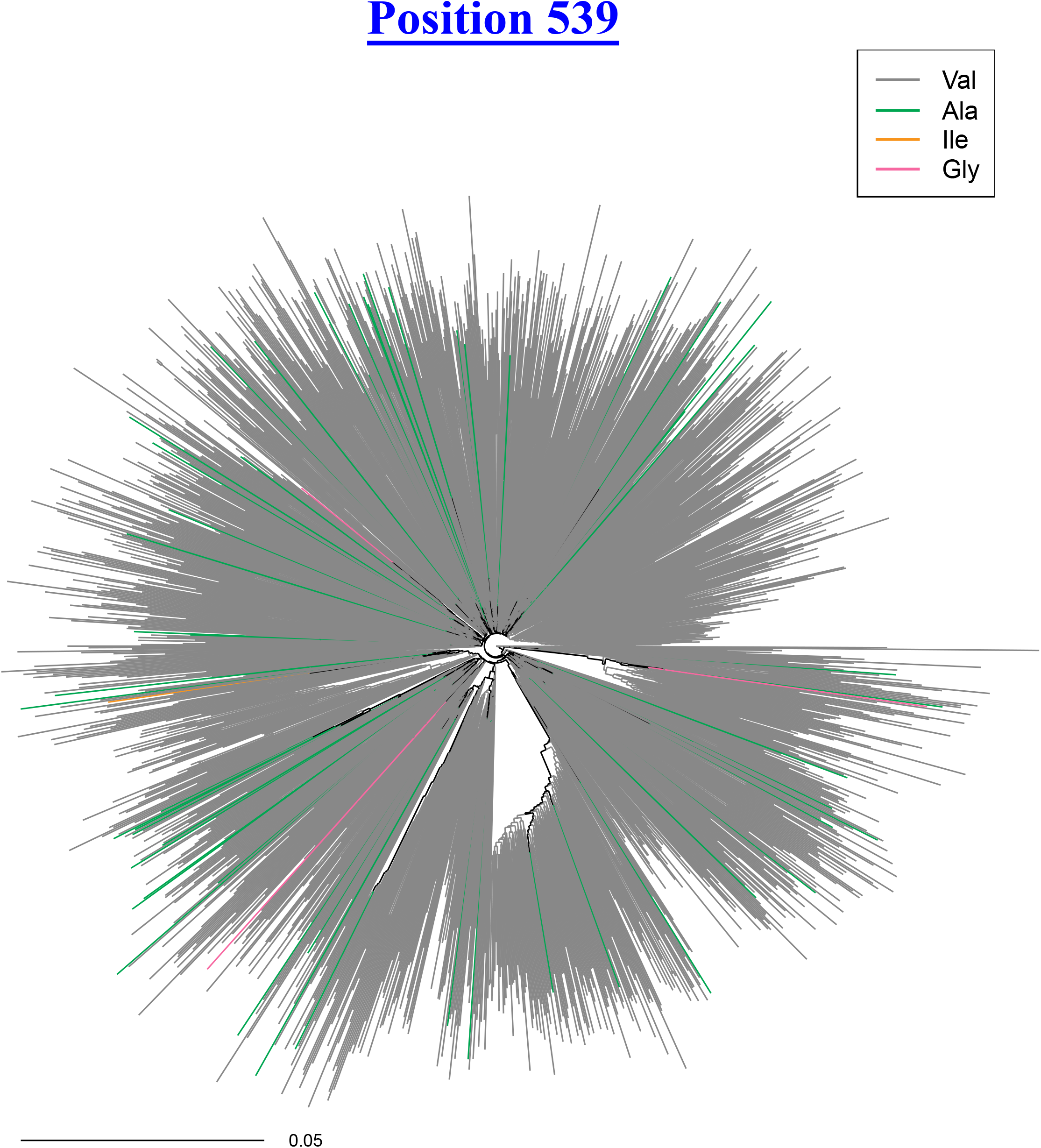
Phylogenetic trees of clade B viruses demonstrating the distribution of amino acids at positions 534, 536 and 539. Nucleotide sequences of HIV-1 *env* from 1,576 distinct individuals were aligned using the HMMER software. Sequences are derived from blood samples collected worldwide between 1979 and 2016. The phylogenetic tree was reconstructed from the sequences using the maximum likelihood method. Isolates are colored by the amino acid they contain at positions 534 **(A)** 536 **(B)** and 539 **(C)**.

